# Adenoviral-Based Vaccine Elicits Robust Systemic and Mucosal Cross-Reactive Responses in African Green Monkeys and Reduces Shedding after SARS-CoV-2 Challenge

**DOI:** 10.1101/2022.12.19.521127

**Authors:** Sarah N. Tedjakusuma, Colin A. Lester, Elena D. Neuhaus, Emery G. Dora, Sean N. Tucker, Becca A. Flitter

## Abstract

As new SARS-CoV-2 variants continue to emerge and impact communities worldwide, efforts to develop next generation vaccines that enhance mucosal immunity would be beneficial for protecting individuals and reducing community transmission. We have developed a non-replicating recombinant adenovirus vector (rAd5) vaccine delivered by mucosal administration engineered to express both a protein antigen and a novel molecular adjuvant in the same cell. Here we describe the immunogenicity of three unique SARS-CoV-2 rAd5 vaccine preclinical candidates and their efficacy following viral challenge in African green monkeys. Animals were prime and boost immunized intranasally twenty-nine days apart with rAd5 vaccine candidates containing viral SARS-CoV-2 spike protein alone or in combination with viral nucleocapsid. Mucosal immunization elicited significant increases in antigen-specific serum antibody responses and functional neutralizing activity against multiple variants of concern. Robust antigen specific mucosal IgA responses were observed after a single administration of rAd5 and generated strong cross-reactive neutralizing antibodies against multiple variants including delta. Importantly, all vaccinated animals exhibited a significant reduction in viral loads and infectious particle shedding in both the nasal passages and lower airways compared to unvaccinated controls following challenge with SARS-CoV-2. These findings demonstrate that mucosal immunization using rAd5 is highly immunogenic, confers protective cross-reactive humoral responses in both the circulation and mucosa, and reduces viral loads and shedding upon challenge with multiple SARS-CoV-2 variants.

## Introduction

While currently approved SARS-CoV-2 vaccines reduce severe disease and hospitalization, these formulations delivered by intramuscular injection do not induce protective immunity at mucosal sites. Delivery of vaccine antigens by intramuscular injection predominantly stimulates immune responses restricted to the peripheral circulation with very weak humoral or cellular responses at mucosal sites where viral entry occurs [1]. Further, as variants of concern (VOC) with increasing fitness continue to emerge, vaccine induced serum responses against SARS-CoV-2 are becoming less effective and more short lived [2–4]. Ideally, next generation SARS-CoV-2 vaccines will stimulate both systemic and mucosal immunity, be cross-reactive against emerging variants, and have the capacity to sustainably reduce community transmission and spread.

Inducing robust IgA responses at mucosal sites is essential for blocking viral entry, impeding attachment, and limiting entry into tissues where the primary infection establishes and spreads. Secretory IgA (sIgA) can be secreted as a multimeric protein and directly bind multiple virus particles, obstructing virus-receptor interaction and subsequent entry into target cells (Reviewed by [5]). Due to increased valency, sIgA has been shown to confer superior neutralizing activity against invading pathogens compared to IgG [6, 7] and prevents respiratory and enteric pathogens from infecting mucosal sites by immune exclusion, steric hindrance, and inhibiting intracellular viral production [8–10]. Further, dimeric sIgA was found to be 15 more neutralizing than serum IgG against SARS-CoV-2 [11]. Therefore, generating potent IgA responses to SARS-CoV-2 antigens through mucosal vaccination may greatly reduce both viral shedding and transmission [12], ultimately impacting the trajectory and subsequent phases of the pandemic.

We have developed an oral vaccine platform that uses a replication-incompetent human adenovirus type-5 vectored vaccine (rAd5) with demonstrated efficacy in humans against a different respiratory virus [13]. The final product is thermostable and specifically formulated to be dried and tableted for easy oral administration and distribution. This temperature stable rAd5 oral vaccine has been administered to over 500 human subjects, has an excellent safety profile, is well tolerated, and generates robust humoral and cellular immune responses against targeted antigens [14–16]. In humans, mucosal responses have been shown to be induced in the intestine and the nose following oral tablet delivery [14, 17]. Furthermore, the potency of this mucosal immunization strategy has been demonstrated in an aerosol preclinical hamster SARS-CoV-2 transmission study, where unvaccinated hamsters were protected from disease when co-housed with hamsters that were challenged with virus after mucosal immunization [18]. These studies show that orally administered rAd5 can enhance mucosal responses and reduce aerosol transmission, providing an effective, potential future strategy for reducing viral disease.

To better understand the immunogenicity of the vaccine candidates and select the best clinical candidate to advance, we evaluated three unique rAd5 SARS-CoV-2 vaccine candidates in African Green Monkeys (AGM). The goal of this study was to select a vaccine candidate that induced cross-protective mucosal and systemic immune responses to a diverse group of SARS-CoV-2 variants and protected against infectious respiratory challenge. Our first clinical candidate contained both the ancestral Wuhan spike and nucleocapsid antigens. Since including nucleocapsid may influence immunogenicity to spike, we selectively evaluated vaccine candidate expressing spike alone and spike plus nucleocapsid vaccine antigens. To determine if immunizing with the homologous matching spike antigen would provide more protective efficacy when new emerging variants arise, we examined a variant-specific vaccine approach against the beta variant. Lastly, as protein-based vaccines are being deployed around the world, a protein subunit vaccine prime followed by mucosal rAd5 adenoviral immunization boost was used as a comparator group. We evaluated a two-dose vaccine series of three rAd5 candidates (and one heterologous immunization regiment) to induce systemic and mucosal immune responses to full length trimerized spike and the receptor binding domain (RBD) from circulating VOC. Here we report that a rAd5 based mucosal vaccine has the capacity to stimulate cross-reactive antibody responses both in circulation and in the respiratory tract and reduce viral loads and shedding in AGMs following a SARS-CoV-2 challenge.

## Results

### Mucosal vaccination elicits strong cross-reactive systemic immunity

To obtain a rapid evaluation of each rAd5 candidate to elicit systemic and mucosal immunity, we chose intranasal administration of liquid rAd5 formulation as an immunization approach. The candidate vaccines evaluated in non-human primates (NHP) utilize replication-incompetent rAd5 recombinant technology designed to deliver vaccine antigens and a molecular dsRNA adjuvant to the same cell (**Fig 1A**). The clinical vaccine candidates ED88, ED90 and ED94 contain codon-optimized transgene sequences of either SARS-CoV-2 spike alone or in conjunction with nucleocapsid (**Fig 1B**). The rAd5 vector ED88 is comprised of two transgenes, Wuhan spike and nucleocapsid under the control of CMV and β-actin promoters, respectively. Whereas ED90 and ED94 rAd5 vectors includes sequences from Wuhan or beta spike, both under the control of a CMV promoter.

**Figure 1:**
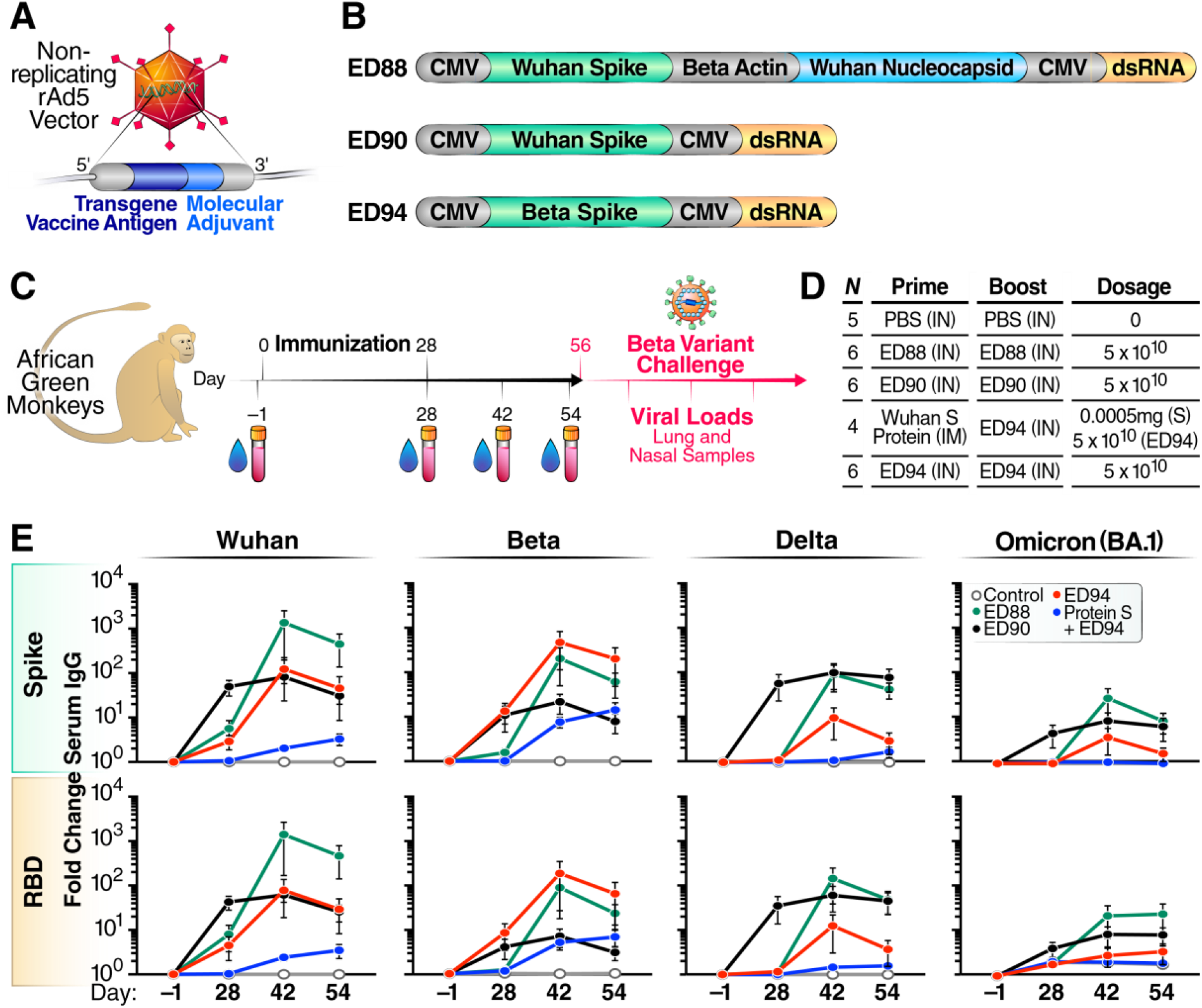
Mucosal immunization with ED88, ED90 and ED94 generates cross reactive serum IgG. **(A)** Illustration of nonreplicating adenovirus type-5 vaccine engineered to deliver both transgene antigen and a dsRNA molecular adjuvant to the same cell. **(B)** Schematic of transgene antigens in rAd5 SARS-CoV-2 vaccines ED88, ED90, and ED94. **(C)** Diagram of study design indicating vaccine administration schedule, serum and nasal swab sample collection, and infectious challenge with SARS-CoV-2 beta variant. **(D)** Vaccines administered during prime and boost immunization and number of animals assigned per group. **(E)** Serum spike specific IgG was quantified by Meso Scale Discovery (MSD) against Wuhan, beta, delta and omicron (BA.1) variants on D-1, D28, D42 and D56. Vehicle control animals denoted (open white circles), ED88 (green circles), ED90 (black circles), ED94 (red circles), or primed with IM delivery of spike protein followed with ED94 boost (blue circles). Data normalized to total IgG in each sample timepoint and expressed as fold change from baseline at D-1, SEM; top row full length trimerized spike, bottom row RBD.

Four groups of AGMs were immunized with a prime boost regimen on day 0 and day 28 by intranasal administration using a mucosal atomizer device. To assess the levels of mucosal and systemic immune responses elicited in each group, nasal secretions and serum were collected on day −1, 28, 42, and 54 post-vaccination (**Fig. 1C).** Three groups of six animals received two intranasal vaccine doses of either ED88, ED90 or ED94 and one comparator group of four animals received an intramuscular injection prime with recombinant spike protein followed by an intranasal boost immunization with ED94 **(Fig. 1D)**. A control group of five animals was intranasally administered PBS in parallel. On day 56, animals were challenged with 1×10^5^ TCID_50_ SARS-CoV-2 beta variant (hCoV-19/South Africa/KRISP-K005325/2020), and serial mucosal samples were collected to determine viral load and shedding.

We first investigated systemic immunogenicity elicited by ED88, ED90 and ED94 mucosal immunization by quantifying serum IgG responses to trimerized spike protein comprising the RBD domains of Wuhan, beta, delta, and omicron (BA.1) using MSD technology (**Fig 1E**). Immunization with ED90 promoted the strongest serum IgG responses with increases over 100-fold against Wuhan and delta and 10-fold increases against beta and omicron variant antigens. Boost administration of ED90 did not result in further increases in serum IgG levels; however, elevated antibody levels were maintained through D54. Interestingly, a stronger cross-reactive serum IgG response against beta, delta, and omicron full-length spike and RBD was detected in animals after receiving two doses of ED88, which encodes the same spike antigen as ED90. Animals immunized with ED94 had strong serum IgG responses to the matched beta spike and RBD proteins after a single dose, which increased following boost vaccination. In contrast, NHPs first vaccinated intramuscularly with recombinant spike protein exhibited very low serum IgG responses, which increased after intranasal boost immunization with ED94 promoting humoral responses against beta variant spike and RBD.

As expected, animals immunized with matched vaccine antigens generated slightly higher IgG responses against the corresponding full-length spike and RBD variant proteins. However, we observed that ED94 vaccinated animals elicited lower cross-reactive serum IgG responses to other variants compared to AGMs immunized with ED90 and ED88. Of note, all vaccinated groups elicited similar serum IgA responses against Wuhan, beta, delta, and omicron spike and RBD proteins, but lower in magnitude compared to circulating IgG. (**Supplemental Fig 1**). These results demonstrate that intranasal administration of matching rAd5 vaccines generated the highest magnitude of IgG and IgA responses to the corresponding spike protein variants, while vaccine candidates ED90 and ED88 promoted the strongest antibody breadth to multiple VOCs.

### Robust cross-reactive nasal IgA is elicited following mucosal vaccination

The production of antigen-specific IgA is important as the first line of defense at mucosal surfaces where the primary viral infection establishes and spreads. To better understand the mucosal humoral response following vaccination with ED88, ED90 and ED94 vaccine constructs, the level of IgA was measured in nasal secretions collected from both nostrils using synthetic absorptive matrix (SAM) devices. (**Fig 2**). Nasal IgA-specific levels against Wuhan, beta, delta and omicron spike or RBD proteins were normalized to total IgA quantified in each respective sample and shown as fold change compared to day −1. After a single administration of ED90, IgA in nasal secretions increased 600-fold to matching Wuhan spike and RBD and over 200-fold to beta, delta, and omicron (BA.1). Following boost administration, the quantity of spike-binding nasal IgA increased in ED90 vaccinated animals to an overall 3000-fold increase from baseline responses. A single administration of ED88 induced a slightly lower increase in nasal IgA specific response to Wuhan, beta, delta, and omicron spike and RBD proteins compared to ED90. However, upon boost administration, ED88 vaccinated animals had similar levels of cross-reactive IgA in nasal secretions compared to ED90 immunized AGM. As predicted, ED94 vaccinated animals generated slightly higher nasal IgA responses to the matching beta spike and RBD proteins after a single dose. Interestingly, upon a second immunization of ED94, cross reactive nasal IgA to other variants was similar in magnitude to ED88 and ED90 prime and boosted animals. The AGM vaccinated with spike protein by intramuscular injection did not produce nasal IgA after 28 days; however, following boost with intranasal ED94 vaccination, spike-specific IgA mucosal responses were observed in the nasal secretions at day 54. These results demonstrate that mucosal immunization with ED88, ED90 and ED94 vaccines induced robust increases in cross-reactive IgA levels to multiple VOC in the nasal mucosa.

**Figure 2.**
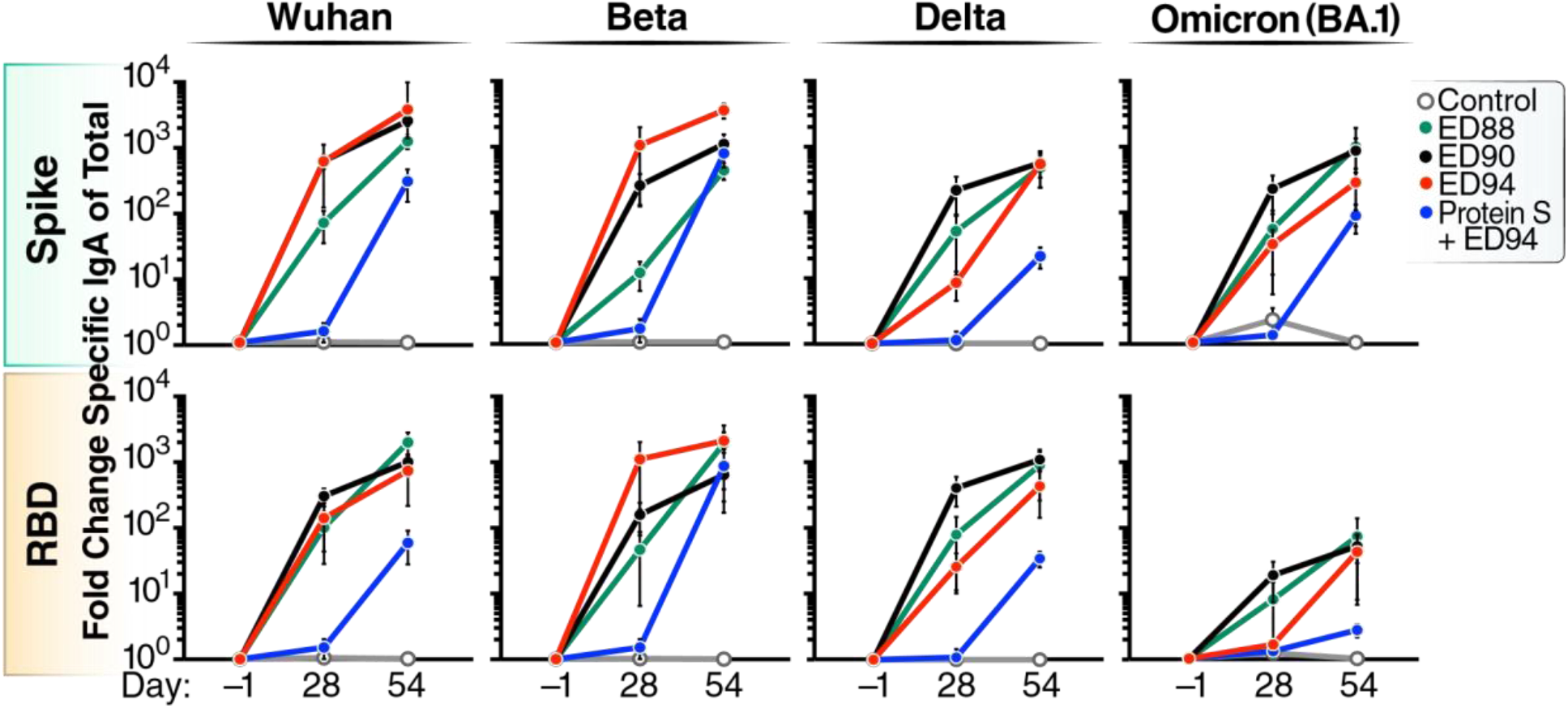
Mucosal immunization stimulates robust cross-reactive nasal IgA to multiple spike protein VOC. Spike specific IgA against Wuhan, beta, delta, and omicron (BA.1) and total IgA were quantified from nasal secretions collected at D-1, D28, and D54 post immunization. Vehicle control animals denoted (open white circles), ED88 (green circles), ED90 (black circles), ED94 (red circles), or primed on D0 with IM delivery of spike protein followed with ED94 boost (blue circles) on D28. Top row full length trimerized spike; bottom row RBD protein. Data normalized to total IgA in each sample timepoint and expressed as fold change from baseline at D-1, SEM.

### Boost immunization enhances neutralizing antibody activity in both the peripheral and mucosal compartments

After observing high amounts of antibodies in the mucosa and serum that bound to spike and RBD proteins, we wanted to ascertain if intranasal rAd5 immunization induced functional serum responses that prevent ACE-2-RBD interactions. We first assessed serum functional activity using a surrogate virus neutralization test (sVNT) which measures the ability of serum and mucosal antibodies to block the interaction between SARS-CoV-2 RBD and the cellular receptor, ACE-2. ED90 vaccination induced strong levels of neutralizing serum antibodies against Wuhan and delta RBD proteins three weeks after prime vaccination (**Fig 3A**). ED88 vaccinated AGM had significantly higher levels of neutralizing serum antibodies against Wuhan, beta, and delta RBD proteins after boost vaccination. Following a single dose of ED94, slightly lower levels of cross-reactive neutralizing serum antibodies against Wuhan, beta, and delta RBD were generated compared to ED88 and ED90 vaccinated AGM. Boost vaccination with ED94 elicited stronger serum neutralizing responses against the homologous beta RBD compared to ED88 and ED90 immunized AGM. Animals vaccinated with purified spike protein by intramuscular injection did not produce serum neutralizing antibodies after 28 days, and the subsequent single boost of intranasal ED94 did not increase functional activity. Surprisingly, while we observed increases in binding antibodies to omicron BA.1 in the serum, these responses were not able to prevent RBD and ACE-2 interactions.

**Figure 3.**
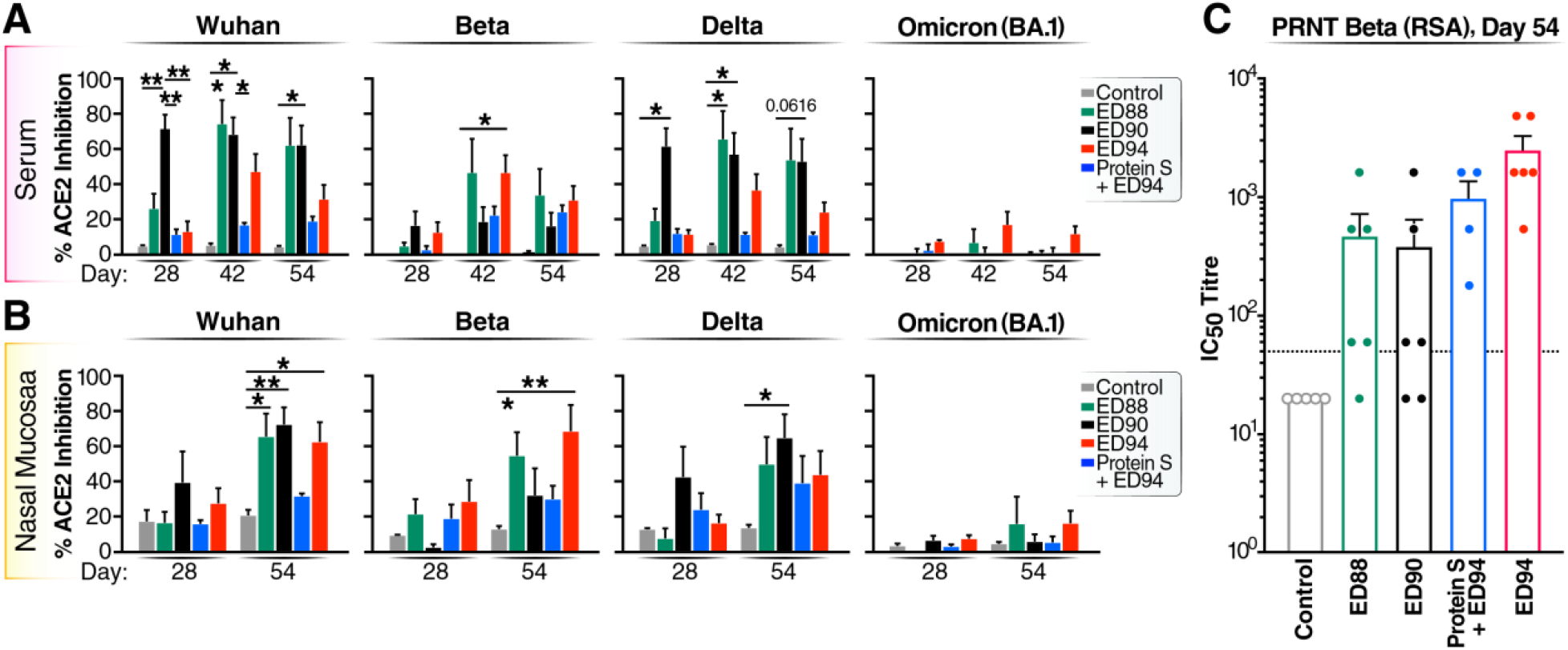
Mucosal vaccination induces neutralizing antibodies in serum and nasal secretions. **(A)** Serum and **(B)** nasal neutralizing antibody activity against the RBD portions of Wuhan, beta, delta and omicron proteins by SVNT. Percent ACE2 inhibition shown on D28, D42 and D54 for serum (top row) and on D28 and D54 for nasal eluants (bottom row). Vehicle control animals (white bars) and vaccinated groups ED88 (green bars), ED90 (black bars), ED94 (red bars), IM delivery of spike protein followed with ED94 boost (blue bars). Data expressed as mean +/− SEM; Two-Way ANOVA with Tukey-Kramer post-test. **(C)** Serum neutralizing IC_50_ antibody titers per animal on D54 by PRNT assay against SARS-CoV-2 beta variant. The dotted line denotes proposed NHP IC_50_ protective serum titer (IC_50_ = 50).

To further confirm our sVNT assay results, a plaque reduction neutralization test (PRNT) was conducted on serum collected on D56, and IC_50_ titers were determined against the SARS-CoV-2 beta variant **(Fig 3C)**. The serum PRNT results verified that animals administered two intranasal doses of ED94 had the strongest serum neutralizing activity to the homologous beta variant with 100% of animals having IC_50_ titers above proposed protective levels in NHP models [19]. Remarkably, 60% of animals primed and boosted with ED88 or ED90 also had protective serum IC_50_ titers, again demonstrating that mucosal immunization with ancestral Wuhan spike induces cross-reactive functional serum humoral responses **(Fig. 3B).** This result confirmed our earlier findings in a Cynomolgus macaque model, that mucosal administration of ED90 generates cross-reactive functional antibody in the serum [20]. Furthermore, the neutralizing serum antibody responses observed were similar to what has been reported with other adenoviral and mRNA vaccines [21, 22].

While neutralizing antibodies in the serum is important for systemic inhibition of the virus and preventing severe disease, these protective responses are restricted to the periphery and unable to limit primary viral infection at the mucosal surfaces. Considering the high levels of antibody binding to spike and RBD VOC observed in the nasal samples, we wanted to determine if these antibody responses also confer functional neutralization activity. To this end, we again employed the sVNT assay to assess neutralization antibody activity intranasally. We found that two intranasal doses of ED88, ED90 or ED94 were required to induce potent neutralizing activity in the nasal mucosa **(Fig 3B)**. Administration of recombinant spike protein by intramuscular injection followed by a single intranasal ED94 vaccination generally did not generate mucosal neutralization activity. Prime immunization with ED88 and ED90 generated the strongest cross-reactive neutralizing antibody response against Wuhan, beta and delta VOC. Two doses of ED94 also generated mucosal neutralizing antibodies to Wuhan and beta, but less so to delta. While we observed strong increases in antibody response to omicron BA.1 spike in the nasal samples from ED88, ED90 and ED94 vaccinated animals, these responses were lower against the RBD and conferred weak neutralizing activity. Overall, these results demonstrated that mucosal administration of ED88, ED90, and ED94 vaccine candidates elicited strong cross-reactive functional neutralizing antibody in both the serum and mucosal compartments.

### Viral replication and shedding is significantly reduced in immunized animals after challenge

Designing vaccines that induce protective mucosal immune responses and limit productive infection and shedding in the upper respiratory tract could have a major impact on the viral disease transmission. To examine whether mucosal immunization reduces viral load following challenge, viral mRNA was measured in the nasal samples and bronchoalveolar lavage fluid (BALF) by qPCR. We used genomic mRNA to quantify total viral transcripts and subgenomic mRNA to measure active viral replication. 24 hours post-challenge, all vaccinated groups had considerably lower genomic and subgenomic viral mRNA detected in the nasal swabs compared to the unvaccinated control group **(Fig. 4A**). Of note, the viral mRNA transcripts continued to decrease over time in nasal passages on D2, D5 and D8 post-challenge in all animals vaccinated with rAd5. Furthermore, on D5, subgenomic transcript levels fell below the limit of detection in animals that received ED94 vaccination, the vaccine that matched the beta challenge strain. In the lower airways, all groups had high viral loads on D2 post-challenge in the BALF which rapidly decreased by 4 logs on D8 **(Fig. 4B)**. Interestingly, animals immunized with purified spike followed by ED94 intranasal vaccination had the lowest levels of viral mRNA in the nasal passages, but the highest level of transcripts in the lower airways. This observation suggests that two doses of intranasal ED94 are required to substantially reduce viral replication in the lower airways. These data demonstrated that mucosal immunization with either matched (ED94) or mismatched (ED88 and ED90) vaccines effectively reduced viral load in both nasal passages and lower airways.

**Figure 4.**
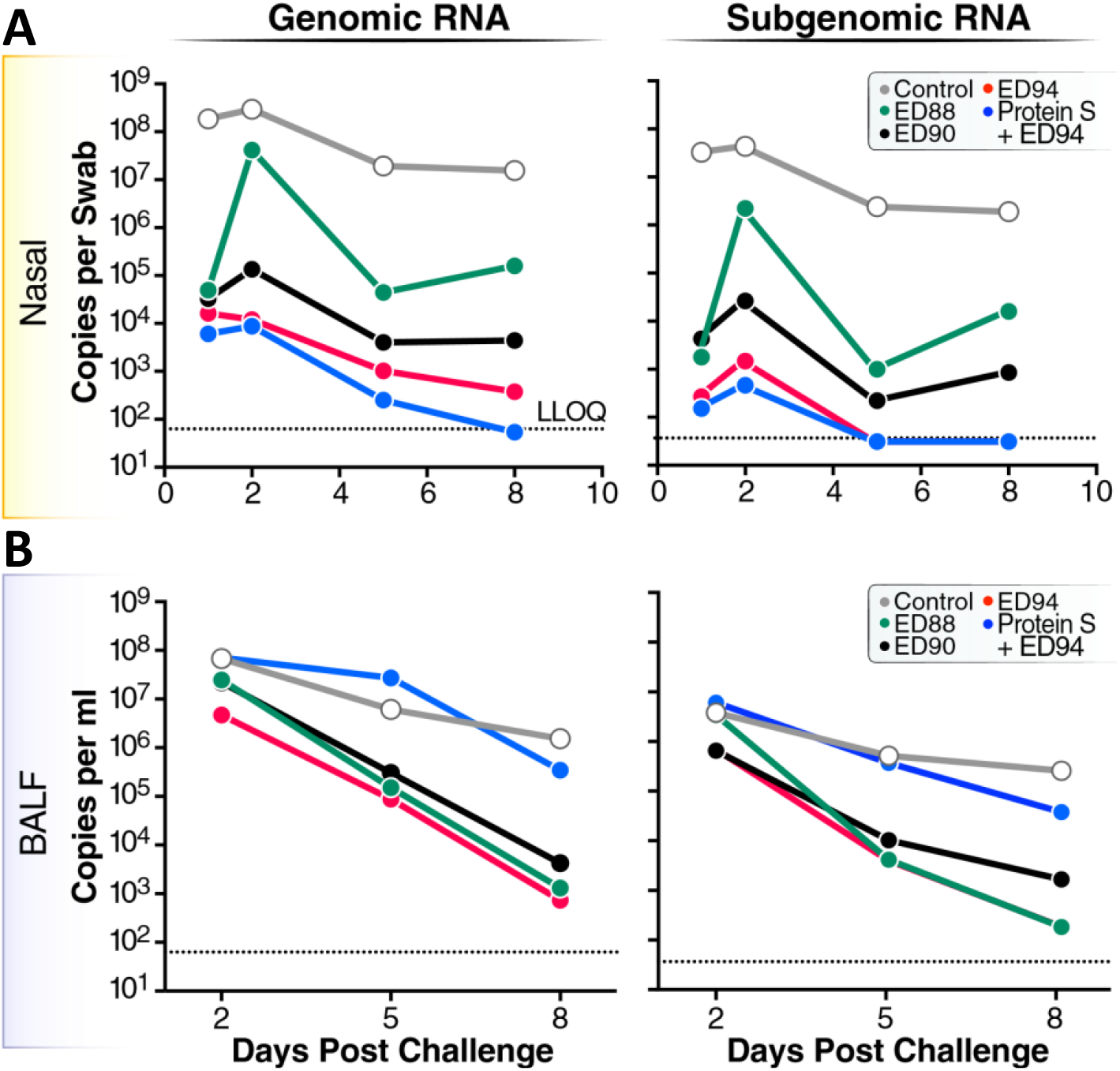
Viral loads are dramatically reduced in nasal mucosa and lower airways of vaccinated animals after challenge with SARS-CoV-2 beta variant. Genomic and subgenomic mRNA was quantified by qPCR from **(A)** nasal swabs on D1, D2, D5, and D8 or **(B)** BALF on D2, D5, and D8 days post-challenge. Vehicle control animals (open circles), vaccinated animals ED88 (green circles), ED90 (black circles), ED94 (red circles), or IM prime of spike protein followed with ED94 boost (blue circles). Data expressed as mean for each group; copies per swab for nasal mucosa and copies per ml for BALF. LLOQ is lower limit of quantitation.

To determine if vaccination with ED88, ED90 and ED94 could reduce infectious viral shedding, mucosal samples were quantified by TCID_50_ from nasal swabs and BALF. High infectious viral shedding, observed in the nasal mucosa in unvaccinated control animals, peaked on D2 post-challenge and was still detectable on D8 **(Fig 5A**). In contrast, infectious virus significantly decreased in all vaccinated animals 24 hours after challenge compared to the control group. Notably, all immunization regimens reduced the amount of infectious viral shedding below lower limits of quantification in most samples. Surprisingly, while the four animals administered spike protein intramuscularly followed by a single dose of ED94 had less mucosal and systemic humoral immunogenicity compared to the other immunized groups, these animals were still protected against the matched challenge beta variant and had low levels of shedding in the nose. In contrast to the nasal mucosa, infectious virus particles detected in the lower airways were only somewhat reduced in immunized animals, with only NHPs vaccinated with two doses of ED94 having significant decreases in shedding **(Fig 5B**). While immunizations with mismatched vaccine antigens were cross-protective in the nasal mucosa, vaccination with ED94 was more effective at reducing viral shedding in the lower airways upon challenge with a matching variant. These results demonstrate that intranasal vaccination with ED88, ED90 or ED94 significantly reduced infectious viral load and shedding in the respiratory tract of AGMs after challenge.

**Figure 5.**
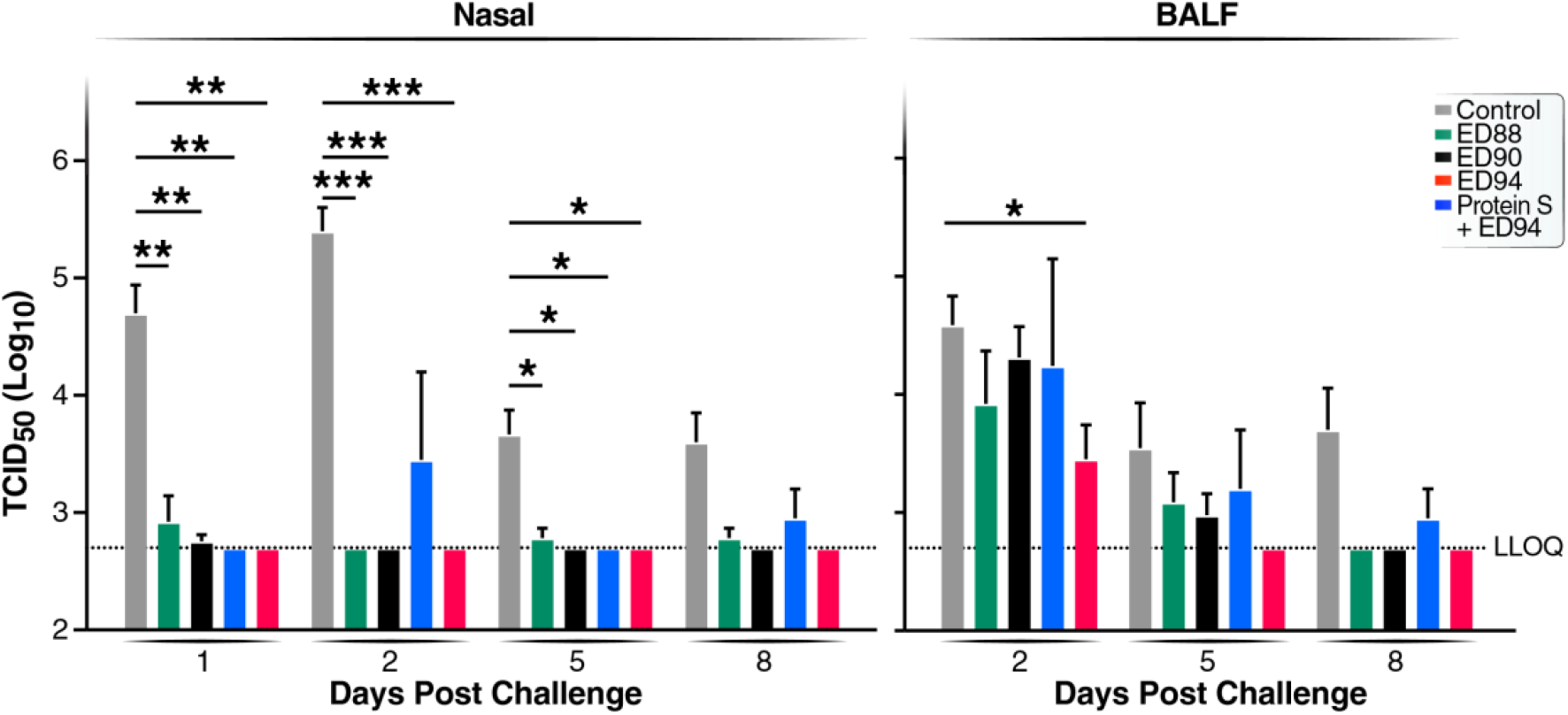
Shedding of SARS-CoV-2 in vaccinated animals is significantly reduced in upper and lower airways. Infectious SARS-CoV-2 beta variant virus shedding was quantified by TCID_50_. Post viral challenge, (**A**) nasal swabs collected on D1, D2, D5, and D8 days and (**B**) BALF on D2, D5 and D8 were plated on Vero cells and TCID_50_ value was calculated using the Read-Muench formula. Vehicle control animals (open bars), immunized groups ED88 (green bars), ED90 (black bars), ED94 (red bars), or IM prime of spike protein followed with ED94 boost (blue circles). LLOQ = 2.7. Data expressed as log10 TCID_50_ /mL, mean +/− SEM.

## Discussion

Here we report rAd5 SARS-CoV-2 vaccines are highly immunogenic in AGMs and elicit cross-reactive functional antibody responses in the circulation and mucosal surfaces, confirming our previous immunogenicity results in Cynomolgus macaques [20]. We observed that all rAd5 vaccine candidates tested were able to generate cross-reactive antibody responses to mismatched antigens. The ED90 vaccine candidate had the highest magnitude and antibody breadth to Wuhan, beta, delta and omicron BA.1 variants, which is similar to our previous findings [20]. Serum neutralization responses following rAd5 vaccination was similar to other published NHP SARS-CoV-2 studies where PRNT responses peaked after two doses [20, 23]. In addition to the strong systemic antibody breadth and functional activity, all three rAd5 vaccine candidates also induced potent mucosal IgA responses with substantial neutralizing activity in the respiratory tract. We observed that two intranasal vaccine doses was the was most effective for inducing potent neutralizing cross-reactive mucosal IgA responses. While both circulating and mucosal antibody responses trended slightly higher to RBD spike proteins when animals were immunized with the matching antigen, sufficient cross-reactive antibody was generated to mismatched antigens that conferred functional protective responses *in vivo* following challenge.

Following SARS-CoV-2 challenge, all rAd5 SARS-CoV-2 vaccines candidates significantly reduced viral load and shedding in the upper respiratory tract demonstrating this immunization approach limits productive viral replication in the mucosa. In comparison to other NHP SARS-CoV-2 challenge studies, we observed significantly lower infectious viral titers in as early as two days post-challenge in the BALF [24]. Further, while cross-reactive humoral and cellular responses have been demonstrated in rhesus macaques challenge models following intramuscular vaccination, viral loads were not measured sequentially, and shedding was not determined [25]. Other adenoviral vaccines formulations administered intranasally in rhesus macaques have been tested in SARS-CoV-2 challenge studies [26]. However, it is unknown if the elicited mucosal antibodies were cross-reactive as responses were only measured against Wuhan. Here our results clearly establish that inducing vaccine-specific cross-reactive mucosal responses can confer protective immunity at sites of viral entry. Mucosal immunity elicited by rAd5 vaccination severely blunted primary infection in the respiratory tract and reduced viral shedding for over 7 days. Therefore, this vaccine approach has the potential to limit the spread of disease by reducing viral shedding and limiting transmission of infectious aerosols.

There were two limitations in this study. First, due to the limited availability of cynomolgus and rhesus macaques, we had to use African green monkeys, which unfortunately develop reduced lung pathology, weight loss or temperature fluctuations after challenge. If we had shortened the time following challenge to assess lung pathology, it is possible that we could have observed more inflammation in the control group. However, our main interest was to determine the duration of viral load and shedding, therefore, we waited one week post challenge to conduct lung histology. While the results were not significantly different, the pathology report indicated inflammatory cells found in the airways trended downwards in the group of AGMs that received two intranasal doses of ED94.

An additional limitation of the study was the use of the nasal passages as the route of vaccine administration. Due to the ineffectiveness of human dose solid forms in monkeys, intranasal delivery was used as a proxy. While fantastic results using intranasal delivery as a method to protect against SARS-CoV-2 in animals have been shown [26, 27], studies in humans have not been as impressive. Intranasal delivery of rAd for COVID vaccination in humans was unsuccessful in two different studies [28, 29], and while an intranasal vaccine was approved in India, not much data has been publicly available. One possibility is that the constant endemic coronavirus infections experienced over a human lifetime may have produced significant cross-reactive sIgA in the upper respiratory tract and hindered the ability of the recombinant spike protein from being recognized by naïve or memory B cells. For example, Flumist, is more effective in young children than in older adults [30]. More consistent results were observed when the adenovirus bypassed the upper respiratory tract and was delivered by aerosol [31] or by oral delivery in the intestine [17].

Interventions that induce antibodies in mucosal sites may be superior at preventing infection and reducing transmission of respiratory pathogens. For example, in a Guinea pig influenza infection study, injected sIgA antibodies that homed to the nasal mucosa were able to prevent transmission from infected animals better than injected IgG antibodies [12]. Influenza viral shedding in humans has also been shown to be significantly lower when individuals were administered an oral vaccine that induced a mucosal sIgA response, compared to individuals immunized with a commercial influenza vaccine, despite the commercial vaccine inducing 10-fold higher serum neutralizing antibody titers [32]. Furthermore, in a study led by Duke university, SARS-CoV-2 aerosol transmission to unvaccinated animals has been demonstrated to be significantly reduced in a vaccine breakthrough study when the rAd was given by oral gavage or intranasally [33]. Animals vaccinated by intranasal, oral, or by injected protein vaccine all had equivalent amounts of virus in the upper respiratory tract 1 day after breakthrough infection, but the *in vivo* infectiveness of the mucosal vaccinated animals was substantially less. Because there was higher sIgA levels in the mucosa, it was presumed that this transmission protection was mediated by nasal sIgA. Therefore, rAd5 vaccines can elicit robust immune responses in multiple pre-clinical models, which suggests that these candidates could also induce strong humoral and mucosal immune responses in human trials.

Currently used vaccines protect against hospitalization and severe illness, but do not appear to be as effective at blocking infection or inhibiting transmission. In a study in Singapore by Chia et al, vaccinated subjects were still capable of being infected by the delta variant at similar rates to unvaccinated subjects, however, the vaccinated subjects cleared the virus slightly faster [34]. In a Danish household study, it was determined that a full-vaccination regiment plus a booster was much more likely to prevent transmission of delta than omicron, with an odds ratio of 3.66 [35]. However, household transmission still occurred 25-32% of the time even with a highly vaccinated population. These studies highlight that the prevention of infection and viral transmission by current vaccine regimens is still a major hurdle towards ameliorating the large social, medical, and economic impact of the SARS-CoV-2 pandemic.

Even when a small portion of the population has been mucosally immunized, it can have a large impact on viral transmission dynamics. In a human study, schools were randomly chosen to offer vaccination with Flumist (an intranasal replicating influenza vaccine) and household transmission rates were compared to similar schools where vaccination was not offered. Despite only a 47% vaccination rate at the participating schools, children in intervention-school households had fewer visits to doctors or clinics for influenza-like illness, and adults in these households had fewer such visits [36]. Further, there were substantially less absentees at intervention schools than unvaccinated schools [36]. Given the increase in nasal sIgA associated with Flumist use in children [37], these results may again suggest how potent mucosal immunity could be.

It will continue to be difficult to protect against the continual wave of newly emerging variants with currently used injected vaccines that induce short-lived circulating antibody with limited breadth. Results in human trials with the mRNA vaccines suggest that the serum antibody responses to each successive variant are reduced in comparison to Wuhan [3], therefore the data presented in this study suggests delivering rAd5 to a mucosal surface may enhance cross-protective responses. The goal for SARS-CoV-2 next-generation vaccines should include not only the prevention of severe disease, but also enhancing mucosal responses and reducing community transmission. Advancing next generation vaccines that are easy to administer, store, and distribute, as well as eliciting immunity both in circulation and in mucosal tissues, is paramount to ensuring equitable and effective global public health responses to future waves of SARS-CoV-2 infections.

## Methods

### Immunization and Study Design

Twenty-seven adult African Green Monkeys (52% Male / 48% Female) were randomly assigned to five groups. Animals were immunized with rAd5 5×10^10^ infectious units (I.U.) on days 0 and 28 by intranasal administration using a mucosal atomization device (MAD) which delivered approximately 0.1 mL vaccine construct per nostril. Three groups were administered either ED88, ED90, or ED94 for both prime and boost, and a fourth group was given an intramuscular injection of purified S protein, NR-52308n (BEI Resources) followed by an intranasal boost of ED94. Serum was collected on day −1 from each animal prior to immunization and on day 28, 42, and 54 post vaccination and pre-challenge (+/− 24 hours). Nasal secretions were collected from right and left nostrils using two synthetic absorptive matrix (SAM) Nasosorption FX-i #NSFL-FXI-15 swabs (Mucosal Diagnostics), on D0, D28, and D54, and were immediately frozen and stored at −80°C.

### SARS-CoV-2 Challenge and Specimen Collection

On day 56, all animals were challenged with SARS-CoV-2 isolate h-CoV-19/South Africa/KRISP-K005325/2020 by administering 5×10^4^/mL TCID_50_ by intranasal and intratracheal route for a total 1×10^5^ TCID_50_ administered to each animal. Post-challenge, animals were monitored daily for any abnormal clinical observations. To quantify viral loads and shedding, nasal swabs (FLOQSwab, Copan Diagnostics) were collected on D1, D2, D5 and D8 post-challenge. Bronchioalveolar lavage fluid (BALF) was collected on D54 pre-challenge and D2, D5 and D8 post challenge. BALF sample cellular debris was removed by centrifugation for 8 minutes at 300-400 g at 4°C, and stored at −80°C.

### Adenoviral vaccine constructs

Adenovirus Type 5 (rAd5) vaccine plasmid constructs were created based on the published DNA sequences of SARS-CoV-2 publicly available as Genbank Accession No. MN90847.3 for the parental Wuhan strain or GISAID Accession No. EPI_ISL_678597 for the beta strain. The published amino acid sequences of the SARS-CoV-2 S or N proteins were used to generate nucleotide sequences codon optimized for expression in human cells. These sequences were cloned into the E1 region of the Ad5 genome contained in a plasmid vector using the same vector backbone used in prior clinical trials for production of oral rAd5 tablets [13, 14]. The rAd5 genome was linearized and transfected into the permissive Expi293F suspension cell-line (Thermo Fisher Scientific). The recombinant vaccines were amplified in Expi293F cells and purified by CsCl density centrifugation as previously described ***(ref).***

### Serum IgG and IgA antibody responses to SARS-CoV-2 variants by MSD

Serum IgG and IgA responses to trimerized S protein and RBD was quantified against Wuhan, alpha, beta, and gamma variants using V-PLEX SARS-CoV-2 panel 7 (Meso Scale Diagnostics, MD). A 4-sector U-PLEX (Meso Scale Diagnostics, MD) assay was developed to measure beta (B.1.351) delta (B.1.617.2) and omicron (BA.1) variant specific serum IgG and IgA. 66nM of biotinylated full-length trimerized delta spike and RBD, omicron spike and RBD (Acro Biosystems, DE) were conjugated to four different linkers and multiplexed on a 4-sector U-PLEX plate according to manufacturer protocols. For both V-PLEX or U-PLEX assays, serum was diluted 1:1000 in 1% ECL Blocking Agent (Cytiva, United Kingdom) in 1X PBS, 0.05% Tween 20, assayed in duplicate, and incubated at room temperature (RT) for two hours shaking at 700 rpm. Plates were washed with 1X PBS, 0.05% Tween 20 and incubated with 1:200 dilution of 200X sulfo-tag anti-IgG or IgA (Meso Scale Diagnostics, MD) for 1 hour shaking at 700 rpm. Plates were developed using MSD Gold Read Buffer and data acquired using the MSD Sector Imager 120 instrument.

### IgA detection in nasal secretions by MSD

Antibodies collected from 3mm synthetic absorptive matrix (SAM) Nasosorption FX-i devices (Mucosal Diagnostics, Midhurst, England) were thawed at 20-22°C for 15 minutes and transferred into Eppendorf tubes containing 300 μL elution buffer (0.05% Tween, 1% BSA, 1X PBS). The absorbent tips were cut and vortexed inside the Eppendorf tubes for 30 seconds and the resulting liquid was transferred into pre-charged elution columns (Costar Spin-X, Corning). The elution columns were spun to remove debris at 16,000 g at 4°C for 20 minutes. The left and right nasal eluants were then pooled, aliquoted, and frozen at −80°C prior to use. To measure IgA in the nasal secretions, eluants were diluted 1:2 and 1:4 in 1% ECL Blocking Agent (Cytiva, United Kingdom) and run on a V-PLEX SARS-CoV-2 (Panel 7) or 4-sector U-PLEX assay (Meso Scale Diagnostics, MD) as described above. Antigen specific nasal IgA was normalized to the total amount of IgA in the corresponding sample. Fold change was calculated by dividing the normalized antigen specific nasal IgA values at each timepoint by the day −1 values.

### Nasal and BALF ACE2 neutralizing antibody response to SARS-CoV-2 variants

Percentages of human-ACE2 blocking antibodies were quantified to measure the surrogate neutralizing antibody response in the nasal cavity and lower airways. Samples were diluted 1:2 and 1:4 in diluent (diluent 100, Meso Scale Diagnostics, MD) and run on a Coronavirus Plate 2 assay (Meso Scale Diagnostics, MD) and a U-Plex assay with SARS-CoV-2 delta, beta, and omicron S1 RBD proteins (Meso Scale Diagnostics, MD). Percent inhibition was calculated by subtracting the ratio of average signal from the sample to average signal from the lowest calibration point. Percent antigen specific inhibition of ACE2 was normalized to total ACE2 inhibition by dividing the antigen specific ACE2 inhibition signal by the total ACE2 inhibition signal.

### Viral load by quantitative Reverse Transcription Polymerase Chain Reaction (qRT-PCR)

Viral load was quantified by measuring the amount of RNA copies per mL using a qRT-PCR assay. Viral RNA was isolated using the Qiagen MinElute virus spin kit (cat. no. 57704). The levels of N gene (subgenomic) mRNA (sgmRNA) were assessed by RT-PCR. Primers and a probe specifically designed to amplify and bind to a region of the N gene messenger RNA from SARS-CoV-2 were used. For amplification, the plates were placed in a sequence detector (Applied Biosystems 7500 Real-Time PCR System) and amplified using the following program: 48°C for 30 minutes, 95°C for 10 minutes followed by 40 cycles of 95°C for 15 seconds, and 1 minute at 55°C. This gave a practical range of 50 to 5×10^8^ RNA copies per swab or mL BAL fluid.

### Primers/probe sequences for RNA copies per mL

2019-nCoV_N1-F :5’-GAC CCC AAA ATC AGC GAA AT-3’

2019-nCoV_N1-R: 5’-TCT GGT TAC TGC CAG TTG AAT CTG-3’

2019-nCoV_N1-P: 5’-FAM-ACC CCG CAT TAC GTT TGG TGG ACC-BHQ1-3’

#### Primers for N gene mRNA

SG-N-F: CGATCTCTTGTAGATCTGTTCTC

SG-N-R: GGTGAACCAAGACGCAGTAT

#### Probe for N gene mRNA

FAM- TAACCAGAATGGAGAACGCAGTGGG -BHQ

### Viral shedding by TCID_50_

Infectious viral load was measured by TCID_50_ assay. Vero TMPRSS2 cells (obtained from the Vaccine Research Center-NIAID) were plated at 25,000 cells/well in DMEM + 10% FBS + Gentamicin and incubated at 37°C, 5.0% CO_2_. After 24 hours, medium was replaced and 20 μL of sample was added to the top row in quadruplicate and diluted 10-fold down the plate. Plates were incubated at 37°C, 5.0% CO_2_ for 4 days, and the cell monolayers were visually inspected for cytopathic effect (CPE). Non-infected wells had a clear confluent cell layer while the infected cells had cell rounding. The TCID_50_ value was calculated using the Read-Muench formula. For samples that had less than 3 CPE positive wells, the TCID_50_ value could not be calculated using the Reed-Muench formula. These samples were assigned a titer of below the limit of detection.

### Serum Neutralizing Antibodies by Plaque Reduction Neutralization Test (PRNT)

PRNT assays were conducted at BioQual as described [19]. Unknown heat inactivated serum samples were serially diluted 3-fold and incubated with 30 pfu/well virus (hCoV-19/South Africa/KRISP-K005325/2020) at 37°C, 5.0% CO_2_ for 1 hour. The serum and virus mixture was then added in duplicate to 175,000 cells/well of Vero E6 cells (ATCC, cat# CRL-1586) in 24-well plates, and incubated at 37°C, 5.0% CO_2_ for 1 hour. 1ml of warm 0.5% methylcellulose media was then added, and the plates were incubated at 37°C, 5% CO_2_ for 3 days. Following the removal of media, the plates were wash with PBS, fixed with cold methanol at −20°C and then stained with 0.2% crystal violet (20% MeOH, 80% dH2O). The stain was removed with dH2O washing, and left to dry for at least 15 min. The plaques in each well were recorded and the IC_50_ and IC90 titers were calculated based on the average number of plaques detected in the virus control wells. A control reference serum with established titer (5,400 IC_50_) was included in each assay as a positive control.

## Competing interests

S.N.Ted., C.A.L., E.G.D., E.D.N., S.N.T. and B.A.F., are employees of Vaxart Inc., and/or have received stock options. EGD and SNT are named as inventors covering a SARS-CoV-2 vaccine. SNT is named as an inventor on patent covering the vaccine platform.

## Supplemental figures

**Supplemental Figure 1.**
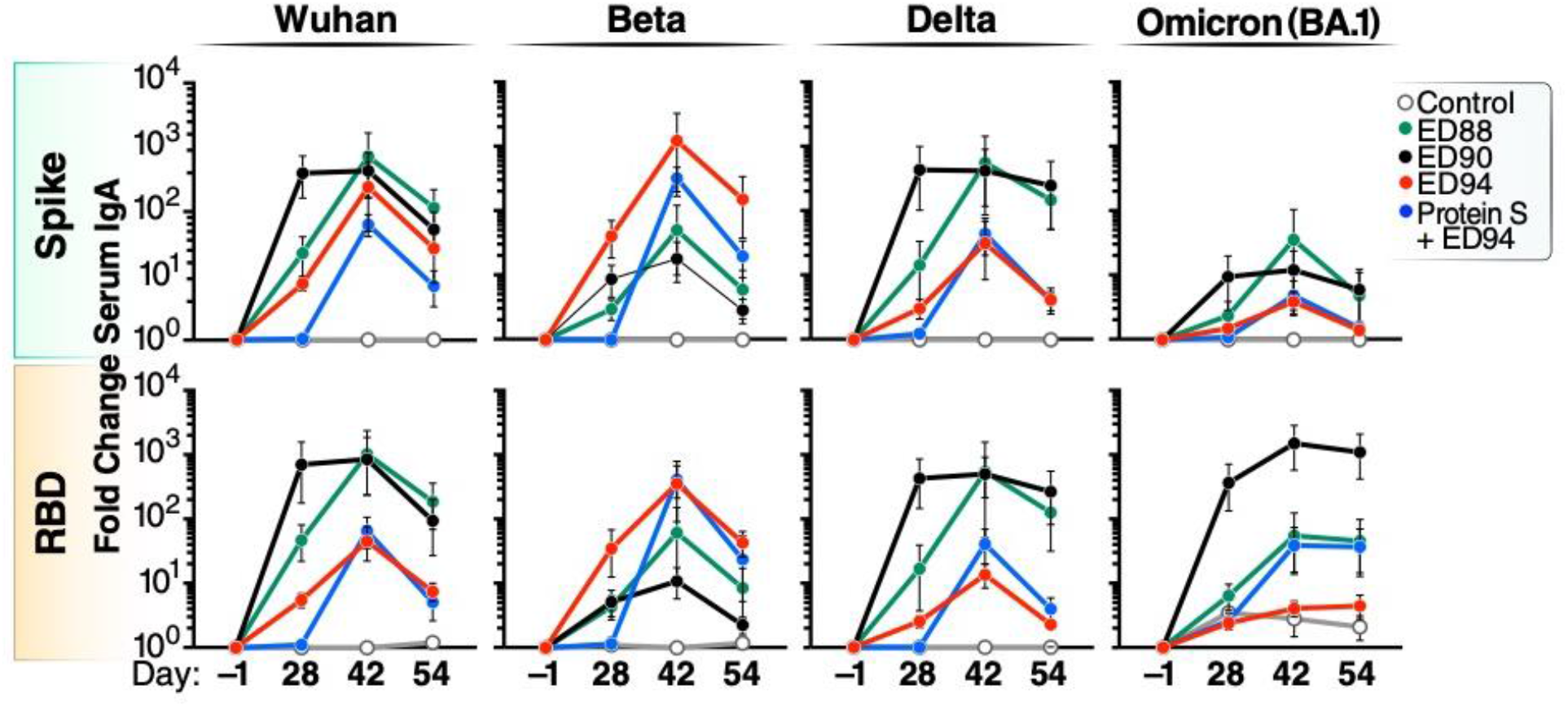
Mucosal immunization generates cross-reactive serum IgA responses. Serum spike specific IgA was quantified by MSD against Wuhan, beta, delta and omicron (BA.1) variants on D-1, D28, D42 and D56. Vehicle control animals denoted (open white circles), ED88 (green circles), ED90 (black circles), ED94 (red circles), or primed with IM delivery of spike protein followed with ED94 boost (blue circles). Data expressed as fold change from baseline at D-1; top row full length trimerized spike, bottom row RBD.

**Supplement Figure 2.**
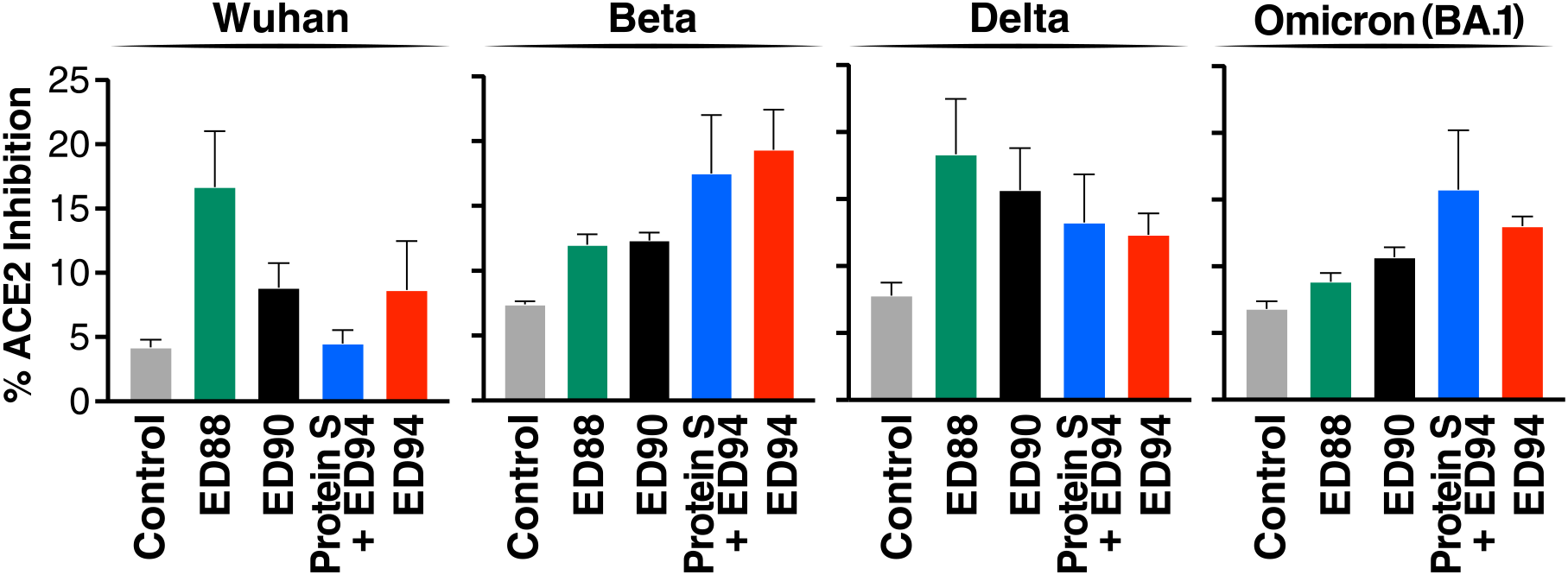
Mucosal immunization enhances neutralizing antibodies in the lower airways. Neutralizing antibody activity against the RBD portions of Wuhan, beta, delta and omicron proteins in BALF by SVNT. Percent ACE2 inhibition shown on D28, D42 and D54 for serum (top row) and on D28 and D54 for nasal eluants (bottom row). Vehicle control animals (white bars) and vaccinated groups ED88 (green bars), ED90 (black bars), ED94 (red bars), IM delivery of spike protein followed with ED94 boost (blue bars). Data expressed as mean +/− SEM.

